# Single-chain permuted proteins for dimerization-based control of protein activity and cellular processes

**DOI:** 10.1101/2025.03.31.645681

**Authors:** Taja Železnik Ramuta, Jaka Snoj, Tina Kos, Ajasja Ljubetič, Iva Hafner Bratkovič, Roman Jerala

**Affiliations:** Department of Synthetic Biology and Immunology, National Institute of Chemistry, Ljubljana, Slovenia; EN-FIST Centre of Excellence, Ljubljana, Slovenia; University of Ljubljana, Faculty of Medicine, Ljubljana, Slovenia; CTGCT, Center for Technologies of the Gene and Cell Therapy, National Institute of Chemistry, Ljubljana, Slovenia

## Abstract

Strategies for detecting and controlling protein interactions play a critical role in gaining insight into molecular mechanisms of biological processes and for the control of cellular processes. Conditional protein reconstitution allows control of the selected protein function based on the proximity, defined by the genetically fused domain pairs, which may be regulated by chemical or biological signals. This typically requires two protein components in a stoichiometric ratio, which increases the complexity and genetic footprint with split segments often being unstable and prone to aggregation. To overcome this limitation, we developed an approach based on a permuted protein reconstitution by conditional dimerization (PROPER). According to this strategy, the N- and C-terminal domains of selected proteins are swapped and a loop is replaced by a short linker that prevents the functionality of a monomeric protein, which reconstitutes only upon di- or oligomerization, controlled by a genetically fused domain that dimerizes by a chemical signal or senses a dimeric target. This design principle was demonstrated on three proteins with diverse functions: a protease, luciferase, and a cytokine. We demonstrate chemically and biologically inducible systems that enable controllable induction of cell death, virus detection, and immune cell stimulation. The PROPER platform expands the chemical biology toolbox, with the benefits of split proteins accomplished by a single component.

## 1 Introduction

Protein interactions play pivotal roles in sustaining all physiological functions, including metabolic pathways, DNA replication, signal transduction, and cell cycle control^1^. Diverse proteins such as receptors, ion channels, proteasomes, and cytoskeletal proteins self-associate to form homodimers or higher-order oligomers to support various biological processes^2,3^. Engineering based on harnessing protein self-association thus presents an important potential for synthetic biology.

Conditional reconstitution systems enable the investigation of protein-protein interactions and protein oligomerization^4^. These systems are often based on splitting a protein into two inactive fragments and coupling the fragments with a module facilitating reassembly. The protein activity in conditional reconstitution systems becomes dependent on specific inducers, such as small molecules^5–8^, light^9–11^, temperature^12^ or proximity binding^13^. The seminal work of Johnsson and Varshavsky^14^, who developed a ubiquitin complementation system that paved the way for linking the specific protein-protein interaction with a split reporter protein reconstitution. A broad array of proteins has been split for a diverse range of applications, extending from addressing universal questions of protein folding and more recently to potential therapeutic applications. Conditional reconstitution systems have been used to study protein-protein interactions (e.g. studies of protein aggregation, identification of cell contacts and synapses)^15–22^ and visualization of cellular processes (e.g. subcellular protein localization, detection of cytosolic peptide delivery)^23–28^, which not only enabled investigation of the function and dynamics of protein-protein interactions in selected biological processes but also supported the identification of molecules that activate or inhibit these interactions. Aside from their role to serve as sensors, conditional reconstitution systems have been implemented in systems that facilitate conditional gene regulation (induced activation or suppression of gene expression using e.g. a split tRNA synthetase, a split transcriptional repressor, INSPIRE system, split Cas9 system)^13,29–31^, genome editing (using e.g. far-red light-activated split-Cas9, split-TALE)^32,33^, and optogenetics (e.g. SPELL system)^9,11^. For therapeutic applications, split locally reconstituted cytokine was shown to decrease toxicity associated with systemic application of cytokines ^34^.

While protein complementation systems present an invaluable tool to monitor or respond to protein dimerization/ oligomerization, their application for the manipulation of cellular functions requires the delivery of both constructs into the same cell. Although a few approaches for the delivery of multiple ORFs within the same plasmid have been recently developed^35–39^, the efficacy of two-component protein complementation systems is limited by variable stoichiometry, high background activity, and a generation of a heterogeneous cell population^40,41^. Additionally, in the case of in vivo application, both constructs need to be stable and resistant to proteolytic digestion^16^. Thus, to avoid these shortcomings new strategies are needed that would allow precise, regulated, and straightforward control of effector protein activity in response to fused protein self-association.

To overcome these limitations, we developed a minimalist single polypeptide chain system, PROPER (*protein permutation and reconstitution*), which enables conditional homodimerization-controlled reconstitution of protein activity. In this strategy, the protein is divided into the N-terminal and C-terminal segments, which are combined in the reverse order, resulting in a biologically inactive permuted protein. The split site is selected so that the monomeric protein is inactive due to a short linker while the dimeric protein regains the activity. Fusion of a permuted protein to a domain that can upon certain conditions dimerize or oligomerize, enables the conditional reconstitution of the protein activity (Figure 1A). The strategy was validated by utilizing several proteins with unrelated functions, including protease, luciferase, and cytokine. Upon dimerization the permuted proteins induced cell death, enabled virus detection, and boosted the activity of immune cells, demonstrating the universality as well as the broad applicability of the PROPER approach.

**Figure 1.**
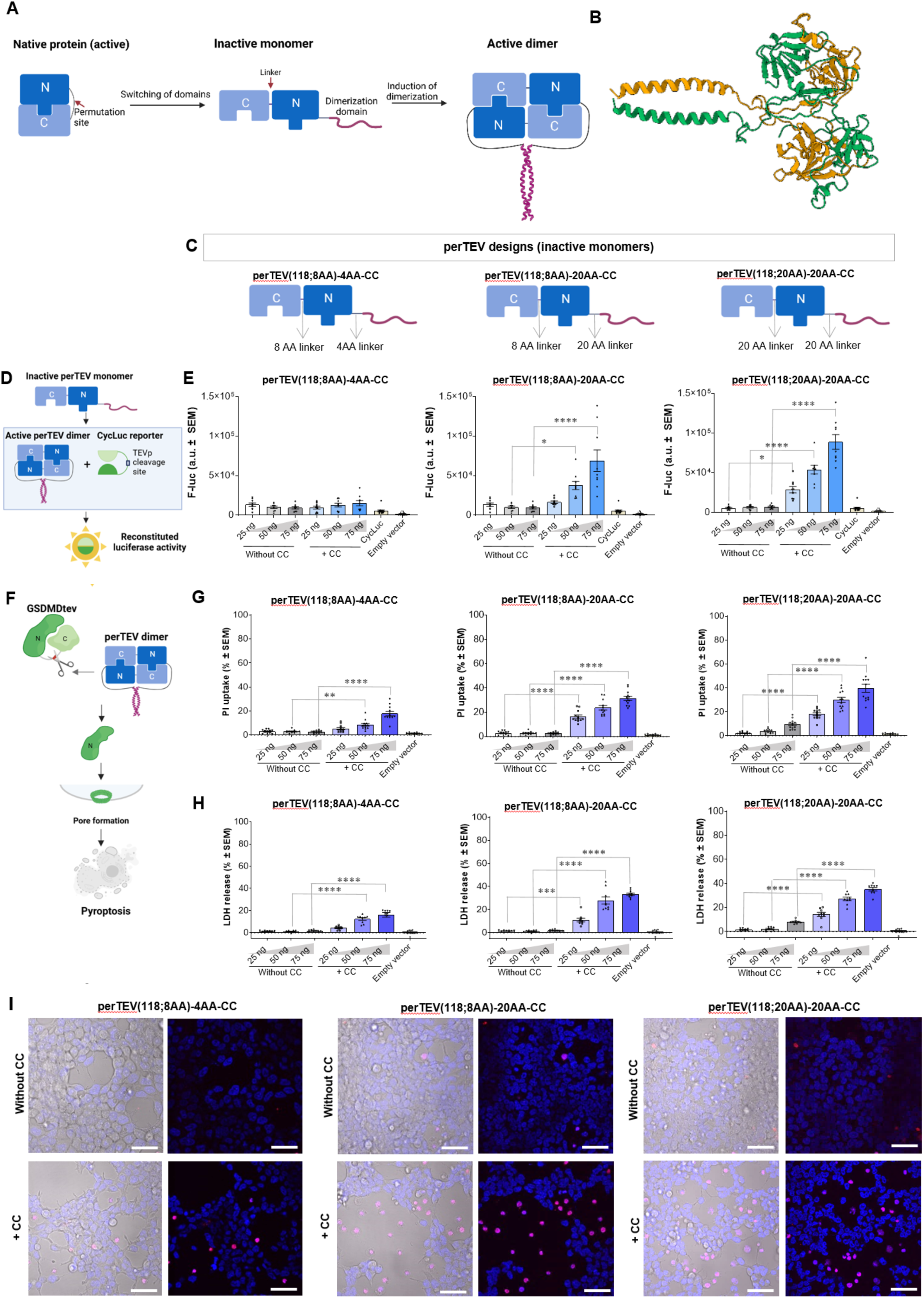
Design and validation of permuted TEV (perTEV) protease. (A) Switching of N and C terminal domains of the original protein sequence and the addition of the linker between domains renders the protein inactive and incapable of reconstitution without additional signals. To establish the PROPER system, the N and C terminal segments are switched, and linkers between protein domains and a dimerization domain are added. Following the signal for dimerization, the perTEVp monomers assemble a dimer resulting in an active enzyme. (B) Protein model of a perTEVp-CC dimer. (C) To engineer the perTEVp the wild-type protein was split after 118AA and three designs were developed, differing in the length of flexible linkers between the protease domains and between the protease and the dimerization domain. (D, E) To determine the activity of perTEVp the HEK293T cells were transiently transfected with the perTEVp and a cyclic luciferase reporter (CycLuc) with a TEV protease cleavage site. Designs without dimerization domains exhibited only minimal activity. (F-H) To determine whether the perTEVp can induce cell death, the HEK293T cells were transiently transfected with the perTEVp and a gasdermin D with a TEV protease cleavage site. To evaluate the extent of induced cell death the propidium iodide (PI) uptake and LDH assay were performed. (I) Induction of cell death by perTEVp designs with dimerization domains as demonstrated by confocal microscopy. Cells exhibiting typical pyroptotic morphology are shown. Dead cells were stained with 7-AAD (red) and nuclei were labeled with Hoechst 33342 (blue). Scale bars 50 μm. Plots show the means ± SEM of at least 9 replicates combined from at least three independent experiments. Conditions were compared using a one-way ANOVA with a Tukey’s multiple comparisons post-hoc test (\*\*\*\**P* < 0.0001; \*\*\**P*<0.001; \*\**P* < 0.01; \**P*<0.05).

## 2 Results

### 2.1 Design principles of dimerization-regulated permuted proteins

The conditionally regulated permuted protein should be inactive as a monomer and active as a homodimer. The design includes the selection of the permutation site, switching the N- and C-terminal domains, and connecting them with an appropriate linker. This can be achieved by two strategies:

1. For proteins whose N- and C-terminus are substantially apart, in the opposing side of the proteins a cyclic permutation with a short linker between the C- and N-terminus prevents folding of a functional monomer. The split site is positioned at one of the surface exposed loops, similar to the preparation of split proteins.
2. For proteins with the N- and C-terminus in proximity, a rigid linker is introduced between the original C- and N-terminus, either as a rigid continuation of a helix or an insertion of a compact protein domain that prevents the permuted protein from folding in a functional monomer.

The additional functional requirement for both cases is that the permuted protein dimer maintains the functional accessibility site of the protein, such as e.g. the binding site of a receptor or access to the catalytic site without a steric hindrance based on dimerization or that there is not a steric clash of a dimer. The linker can also increase the solubility of the monomer, additionally stabilize the dimer and a folded protein domain could act as a rigid linker that could sterically hinder the reconstitution of a monomer. This design resembles to some extent swapped dimers, with the distinction that the dimers should not be formed spontaneously but only upon forced dimerization.

The permuted monomers and dimers split at different sites and with diverse linkers were tested by AlphaFold2 to identify the appropriate candidates. Likely due to the native protein MSA-imposed bias the AlphaFold often predicts the retained monomeric fold of the parent protein, with larger distortions in the loop, and low plDDT values which, as the experimental results show, are likely nonfunctional, while the predicted dimeric structure exhibited more favorable scores.

### 2.2 The design of dimerization-regulated protease

To test the concept of the dimerization-regulated permuted protein design (Figure 1A) we selected tobacco etch virus protease (TEVp). TEVp is a widely used protease in biotechnology due to its highly specific activity at a broad pH and temperature range and no observable toxicity in prokaryotic and eukaryotic cells^51^. To establish the permuted system, we designed several versions of permuted TEV protease (perTEVp) with permissive split sites identified at amino acid positions 70, 118, or 191 (Supplemental Figure 1A), that fulfill the above-mentioned rules. To evaluate if the reconstruction of the permuted protein resulted in a functional protease upon dimerization, it was genetically fused with the GCN homodimerizing coiled coil-forming segment (CC). A model of dimeric protease permuted at site 118 is shown in Fig. 1B. The N- and C-C-termini of the TEV protease were connected by flexible linkers encompassing 8, 12, and 20 AAs. Similarly, the linkers connecting the perTEV protease and the dimerization domain were 4, 10, or 20 AA long (Fig, 1C).

The activity of the designs was tested in transiently transfected human embryonic kidney 293T (HEK293T) cells with the perTEV protease variants and a cyclic firefly luciferase (CycLuc) substrate reporter comprising a TEV cleavage site (Figure 1D, E, Supplemental Figure 1B). The majority of perTEVp designs with splitting site 118 were capable of CycLuc cleavage when fused to a dimerization domain, while in its absence they remained inactive (Figure 1E, Supplemental Figure 1B). On the other hand, the perTEV protease split at position 70 was inactive and the design split at site 191 was constitutively active regardless of the presence of the dimerization domain (Supplemental Figure 1B). The activity between the designs varied in dependence on the length of the linkers. The highest fold induction was achieved by the design with the longest linkers between the protease and the dimerization domain, i.e. perTEVp (118; 8AA)-20AA-CC and perTEVp (118; 20AA)-20AA-CC. The perTEV proteases reached ∼ 50% of the activity of the wild-type TEVp (Figure 1E, Supplemental Figure 1F).

Proteases play a role in diverse cellular functions including as the key executioners of cell death. We were interested in whether the perTEVp designs could be used to trigger an immunogenic cell death such as pyroptosis. In mammalian cells, proteases cleave members of the gasdermin family, licensing the N-terminal cleavage product to assemble the pores in the membranes^52–55^. For this purpose, we used a pyroptosis effector gasdermin D in which the caspase-1 cleavage site has been exchanged for the TEVp cleavage site (GSDMDtev)^56^. Using transient transfection, the GSDMDtev was introduced into the cells together with a perTEV protease (with or without the dimerization domain), and the level of induced cell death was evaluated by measuring a membrane-impermeable dye propidium iodide uptake (an indicator of the GSDMD pore formation), lactate dehydrogenase release (indicator of plasma membrane rupture) and confocal microscopy (Figure 1F-I). The experiments were performed on HEK293T cells which do not express the endogenous gasdermin D and as such provide a clear readout of the synthetically induced cell death. Our results show that all perTEV proteases (split at site 118 AA) fused to the dimerization domain cleaved GSDMDtev resulting in cell death. Moreover, we observed the characteristic ballooning morphology of pyroptotic cells^55^ in cells transfected with an active dimeric perTEV protease (Figure 1I, Supplemental Figure 1E). In accordance with the results using the CycLuc reporter, the perTEVp designs with splits at sites 70 AA and 191 AA were unable to selectively induce pyroptosis regardless of the presence or absence of the dimerization domain (Supplemental Figure 1C-E). The extent of induced cell death correlated with the results on cyclic luciferase reporter. The constructs without the dimerization domain induced cell death in a negligible fraction of cells, comparable to the empty vector (Figure 1E-I, Supplemental Figure 1B-E).

Together, these results demonstrate the successful engineering of a permuted protease, where the split site governs the activity of the permuted protein, which can be further fine-tuned by the linker length. Longer linkers most likely exhibited the least steric hindrance for perTEV dimerization mediated by a dimerization domain.

### 2.3 Chemical and biological regulation of permuted TEV protease activity triggering cell death

The proximity is used to regulate biochemical processes and relay information within and between cells^57^. Chemical inducers of dimerization have been used to regulate signaling pathways and guide cellular processes in animal models^58–60^ and even in therapy^61^, indicating that induced proximity could be used not only to study protein-protein interactions but also for regulation. Chemical (e.g. rapamycin analogs) and biological inducers can be used to achieve the dimerization to regulate the desired signaling, transcription, or localization^62,63^.

To test whether we can use different cues to induce reconstitution of permuted TEV protease and direct its activity, we used the best-performing perTEVp (118AA; 8AA)-20AA design to construct a system that is inducible by chemical or biological signals. To demonstrate that chemically inducible dimerization domains can be applied, permuted TEV proteases were initially fused to FKBP or FRB domains were prepared, which in the presence of rapamycin forms the FKBP-rapamycin-FRB ternary complex. By transiently transfecting the HEK293T cells with a CycLuc reporter and perTEVp-FKBP/FRB constructs, we show that FKBP/FRB heterodimerization system is capable of inducing reconstitution of the permuted TEV protease following the treatment with rapamycin and inducing cleavage of the CycLuc reporter or the pyroptosis effector GSDMDtev (Supplemental Figure 2A-C). As this system lacks the advantage of a single component-mediated regulation, we next designed a chemically inducible system based on a single protein chain, where the perTEV protease was fused to the DmrB homodimerization domain that is inducible by a small molecule, AP20187 homodimerizer ligand (43) (Figure 2A). We demonstrated that the perTEV protease activity is indeed successfully reconstituted by the treatment with the small molecule, while in its absence there was only a minimal level of activity (Figure 2A). Then we tested whether this variant can be used to induce cell pyroptosis as described above. We demonstrated that the B/B homodimerizer successfully induced cell death in a concentration-dependent manner (Figure 2B, C). Moreover, there was no spontaneous reconstitution of the protein as the non-dimerized constructs induced a comparable level of cell death as the non-cleaved GSDMD alone.

**Figure 2.**
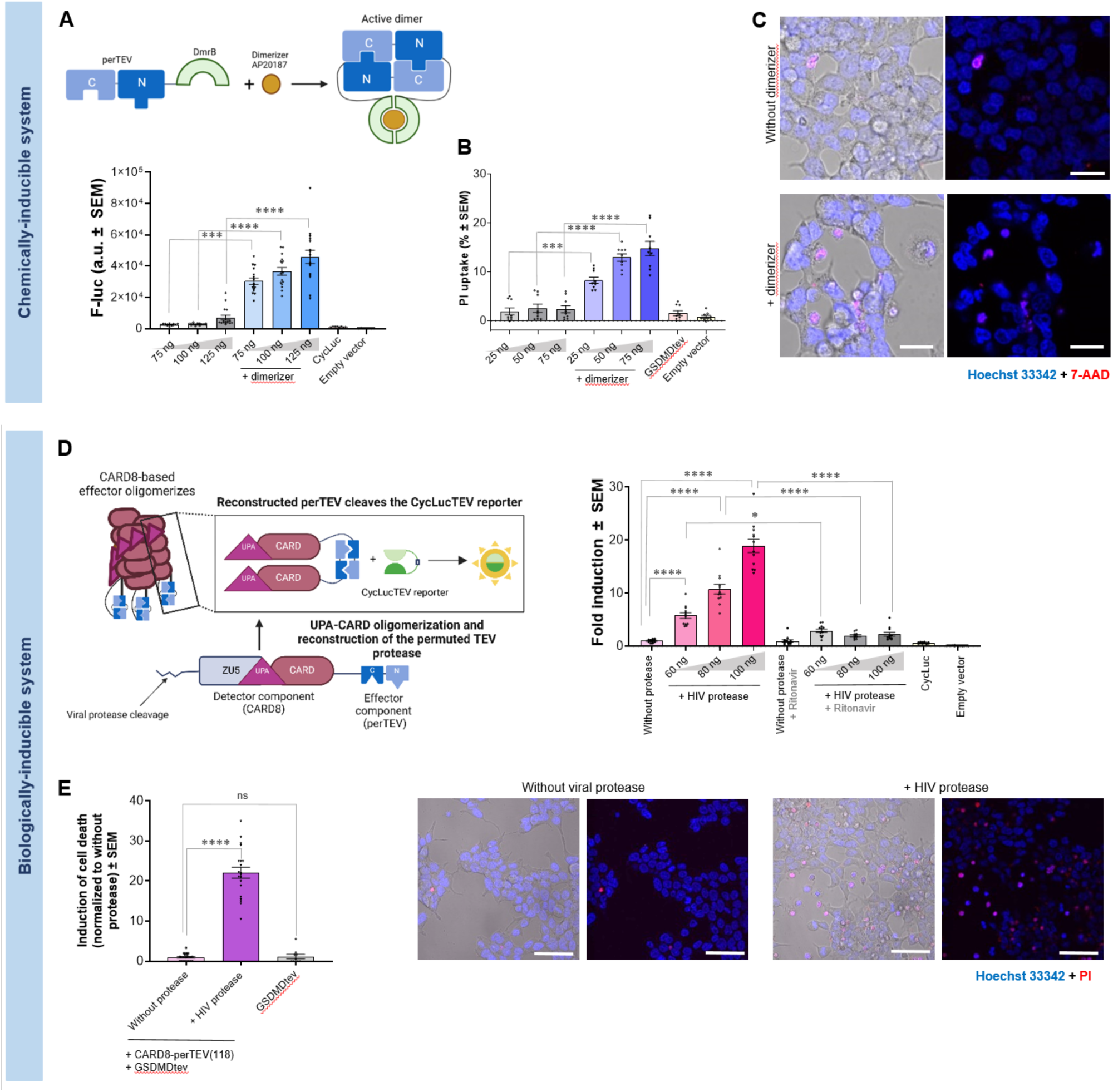
Chemical and biological signals successfully induce reconstitution of perTEV protease. (A) To develop a chemically inducible perTEVp design, the perTEVp with a split at site 118AA was fused to the DmrB homodimerization domain. Following the addition of the small molecule (AP20187) the perTEVp reconstituted and cleaved the CycLuc reporter with a TEV cleavage site. In the absence of the small molecule, the perTEVp-DmrB exhibited only minimal activity. (B, C) The perTEVp-DmrB induced cell death in HEK293T cells transiently co-transfected with GSDMDtev when induced with a small molecule as demonstrated with the PI uptake. Confocal microscopy images show pyroptotic cells exhibiting typical pyroptotic morphology. Dead cells were stained with 7-AAD (red) and nuclei were labeled with Hoechst 33342 (blue). (D) To develop a biologically inducible system the perTEVp(118; 8AA) was fused to the CARD8. As the CARD8 was previously genetically modified it carried the NS3 protease and HIV protease cleavage sites. Thus, the CARD8 served as a detector component and the perTEVp served as the effector component. Following viral cleavage by HIV protease the CARD8-based effector oligomerized and that resulted in reconstruction of the perTEVp which then cleaved the CycLuc reporter in a dose-dependent manner. The addition of the HIV protease inhibitor (Ritonavir) prevented the oligomerization of CARD8-perTEVp. (E) The CARD8-perTEVp successfully induced cell death in the presence of a viral protease in HEK293T cells co-transfected with GSDMDtev. Confocal microscopy images show pyroptotic cells exhibiting typical pyroptotic morphology. Dead cells were stained with PI (red) and nuclei were labeled with Hoechst 33342 (blue). Scale bars (C) 10 μm, (E) 50 μm. Plots show the means ± SEM of at least 9 replicates combined from at least three independent experiments. Conditions were compared using a one-way ANOVA with a Tukey’s multiple comparisons post-hoc test (\*\*\*\**P* < 0.0001; \*\*\**P*<0.001; \*\**P* < 0.01; \**P*<0.05).

To further display the applicability of permuted protein design for integration into natural signaling pathways and their rewiring, we introduced perTEVp as a C-terminal fusion of CARD8 protein (caspase recruitment domain family member 8). Upon viral infection, CARD8 is cleaved by viral proteases, e.g. HIV^64^, enteroviruses^65^, corona- and picorna^66^ viruses. The cleavage at the N-terminus leads to destabilization and proteasomal degradation of the N-terminal part of CARD8 releasing the UPA-CARD segment that polymerizes and forms an inflammasome, leading to caspase-1-mediated GSDMD cleavage and pyroptosis^64^. To demonstrate sensing of viral proteases, we introduced the hepatitis C virus NS3 protease cleavage site adjacent to the inherent HIV protease cleavage site in the N-terminal segment of CARD8 and genetically fused CARD8 to the perTEVp. Upon co-transfection, cleavage of the CARD8-perTEV sensor by the HIV or NS3 proteases led to oligomerization of the CARD8 and subsequent reconstitution of the perTEVp which cleaved the CycLuc reporter. Our results demonstrate that the system enables the detection of protease activity and its response is dose-dependent (Figure 2D, Supplementary Figure 2D). Moreover, the detected signal can be suppressed with the HIV protease inhibitor Ritonavir, demonstrating the specificity of the sensor for detecting the protease activity. This further demonstrates that the permuted proteins are not reconstituted only by dimerization but also by oligomerization either through the reconstitution of several dimers or a chain of oligomers that can reconstitute the functionality.

Similarly, when HEK293T cells were transiently transfected with the CARD8-perTEVp and GSDMDtev constructs, cell death was observed when cells were co-transfected with the NS3 and HIV proteases (Figure 2E, Supplemental Figure 2E, F). Such systems could be used as viral sensors or safety switches that eliminate the cells upon infection with the selected virus.

To demonstrate the functionality in another cell type, several systems were tested in the cancer HeLa cell line, including GSDMDtev and perTEVp(118; 8AA)-20AA (with or without the dimerization domain; CC), perTEVp-DmrB (with or without adding the AP20187 dimerizer) and CARD8-perTEVp (with or without the NS3 protease). All three different systems successfully reconstituted perTEVp with subsequent cleavage of GSDMDtev, leading to cell death in case of induced dimerization/oligomerization, while the cells remain intact in the absence of the activating signal (Supplemental Figure 2G).

We demonstrated that chemically or biologically inducible self-association systems facilitate external control over perTEVp that can serve as a mutation-agnostic sensor of viral protease activity or facilitate selective, signal-dependent cell death.

### 2.4 Permuted nanoluciferase as a sensor of viral oligomeric proteins

Sensitive and efficient high-throughput screening methods are crucial in diagnostics, characterization of viral isolates, and drug development. While (RT)-qPCR-based methods are very sensitive and specific, they typically require an extended time and indicate only the presence or absence of a viral genome or its traces^67^. Rapid antigen tests, on the other hand, demonstrated their use for rapid point-of-care detection of infection despite their lower sensitivity. We aimed to develop a gain-of-signal, luciferase-based assay, that would allow monitoring of the viral presence and receptor binding.

First, we permuted a nanoluciferase (perNanoLuc), which is characterized by enhanced stability and smaller size^68^. The nanoluciferase was permuted at position 158, selected and modeled according to the above-stated rules, and fused to the target protein domains, for detection of viral proteins through two approaches. To develop a sensor that detects the activity of viral proteases, the perNanoLuc was fused to the CARD8 protein with introduced hepatitis C virus NS3 protease cleavage site adjacent to the inherent HIV protease cleavage site in the N-terminal segment of CARD8 (Figure 3A). As a proof of concept, we transiently transfected HEK293T cells with plasmids encoding CARD8-perNanoLuc sensor and NS3 or HIV proteases to simulate viral infection. By measuring bioluminescence, we demonstrated that the viral protease cleavage triggered the CARD8-perNanoLuc sensor oligomerization and reconstruction of perNanoLuc activity. The response of the sensor correlated to the amount of the transfected viral protease encoding plasmid (Figure 3B, C). As viral proteases are crucial for virus replication such detection systems that are based on a viral protease function cannot be overridden by viral protease mutations like antibodies or viral inhibitors.

**Figure 3.**
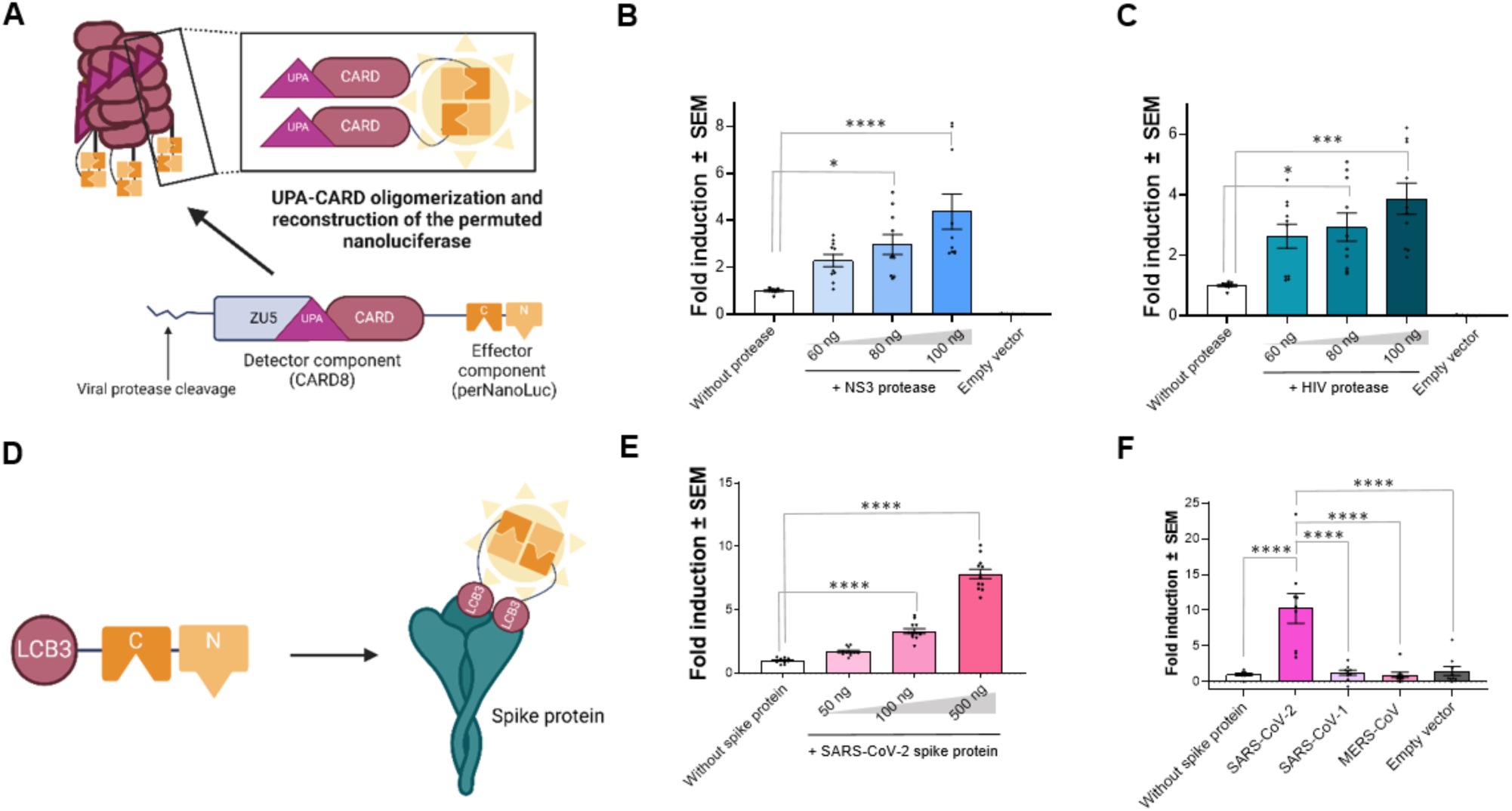
Permuted nanoluciferase as a sensor of viral proteins. (A) To develop a biologically inducible system the permuted nanoluciferase (perNanoLuc) was fused to the CARD8 carrying the NS3 and HIV protease cleavage sites. (B, C) To test whether the CARD8-perNanoLuc sensor successfully detects the viral activity of NS3 and HIV proteases, we transiently transfected the HEK293T cells with the sensor, CycLuc reporter, and NS3 or HIV protease. Following viral cleavage the CARD8-based effector oligomerized and that led to the reconstruction of the perNanoLuc which then cleaved the substrate, resulting in bioluminescence. (D) Scheme of the perNanoLuc-LCB3 sensor for detection of the SARS-CoV-2 spike protein. The perNanoLuc was fused to the LCB3 minibinder that targets the SARS-CoV-2 receptor binding domain. (E) To test the sensor activity of the perNanoLuc-LCB3 sensor and SARS-CoV-2 spike protein, the proteins were first isolated, and in vitro, testing showed that the sensor successfully detected the presence of the SARS-CoV-2 spike protein. The response was dose-dependent. (F) To evaluate the specificity of the perNanoLuc-LCB3 sensor the HEK293T cells were transiently transfected with SARS-CoV-2 / SARS-CoV-1 / MERS-CoV spike proteins and 48 hours post transfection the perNanoLuc-LCB3 sensor was added. The sensor successfully recognized only the SARS-CoV-2, demonstrating high specificity. Plots show the means ± SEM of at least 9 replicates combined from at least three independent experiments. Conditions were compared using a one-way ANOVA with a Tukey’s multiple comparisons post-hoc test (\*\*\*\**P* < 0.0001; \*\*\**P*<0.001; \*\**P* < 0.01; \**P*<0.05).

As an alternative, we tested another principle of a viral sensor for direct detection of viral particles by targeting the surface protein, SARS-CoV-2 spike protein. This sensor design comprises the perNanoLuc fused to the LCB3 minibinder protein, which binds to the SARS-CoV-2 receptor binding domain^69^ (Figure 3D). Spike, similar to most other viral fusion proteins, is a constitutively trimeric protein, and therefore the natural oligomeric state can be used to reconstitute the nanoluciferase activity. As a proof of concept, we isolated the SARS-CoV-2 spike protein and the perNanoLuc-LCB3 sensor (Supplemental Figure 3) and tested binding and sensing in vitro. Our results show that the addition of the trimeric SARS-CoV-2 spike protein^70^ enabled dose-dependent reconstitution of the perNanoLuc activity (Figure 3E). Next, to evaluate the specificity of the perNanoLuc-LCB3 sensor, we transiently transfected the HEK293T cells with the SARS-CoV-2/ SARS-CoV-1/ MERS-CoV spike proteins, which mimics viral infection and added the recombinant perNanoLuc-LCB3 sensor. Only the presence of the SARS-CoV-2 spike protein resulted in the reconstruction of the perNanoLuc, while for SARS-CoV-1 and MERS-CoV the bioluminescence was similar to that of the sensor in combination with cells not expressing spike proteins (Figure 3F), demonstrating the specificity of the sensor.

By developing a permuted version of small and stable NanoLuciferase we showcased the potential of perNanoLuc to be used as a biosensor for the detection of viral or other oligomeric components.

### 2.5 Permuted interleukin 15 enables small molecule-inducible stimulation of natural killer cells

Several strategies have been used to improve the antitumor activity and persistence of NK cells in the tumor microenvironment, the most common being the application of exogenous interleukin 2 (IL-2) or IL-15 to promote the proliferation and cytotoxic activity of NK cells ^71–74^. However, the systemic administration of IL-2 and IL-15 is associated with toxicity^75–78^ and the IL-2 may additionally promote the expansion of regulatory T cells that limit the anticancer activity of NK and other cytotoxic cells^79^. To improve specificity, a split IL-2/IL-15 mimetic has been designed previously that restricts the activation of the cytokine to cells that express tumor antigen or immune checkpoint but this design requires the delivery of two polypeptide components^34^. We aimed to develop a permuted cytokine, to decrease the number of required components to a single protein and to demonstrate that it could promote the anti-tumor activity of NK cells that can be regulated by external small molecules as a safety regulator.

In case of IL-15 N- and C-terminus are in close proximity and to engineer the permuted dimerization-regulated cytokine, IL-15 (perIL-15), the first helix IL-15 of was moved to the C-terminus of the sequence and a single alpha helix was inserted as a rigid linker (Figure 4A) to hinder the functionality of a permuted monomer as an example of the second type of permuted protein generation, as described above.

**Figure 4.**
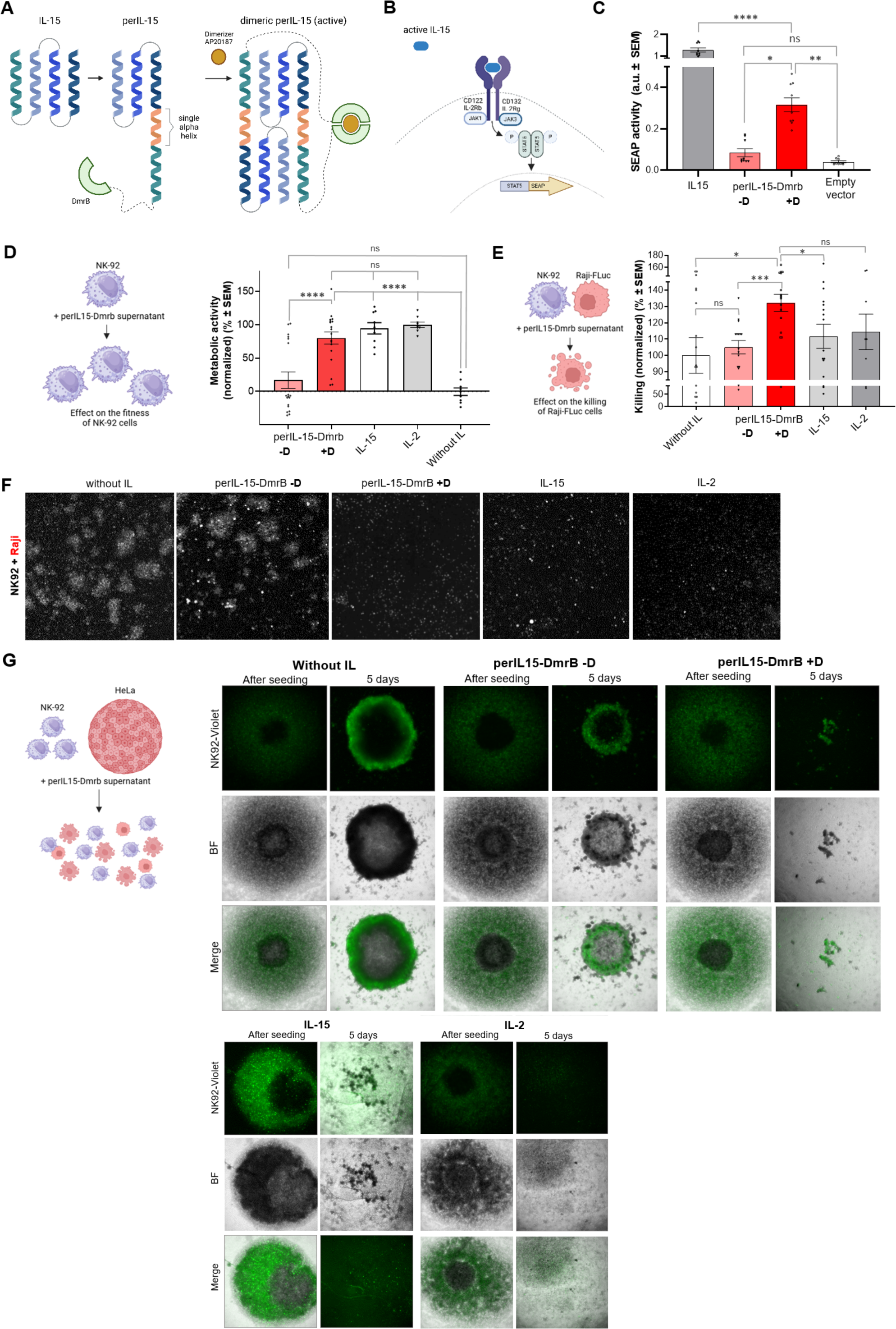
Permuted IL-15 stimulates immune cells. (A) To design the permuted IL-15 (perIL-15) the order of the four alpha helices comprising the IL-15 was switched as the first helix was moved to the end of the sequence and a rigid single alpha helix was inserted between the rest of the helices and the first helix. This construct was then fused to the dimerization domain. (B) To test the activity of the perIL-15 the HEK-Blue CD122/132 reporter cell line was used. Binding of the IL-15 triggered the signaling cascade resulting in the activation of STAT5, leading to the production of secreted embryonic alkaline phosphatase (SEAP). (C) A chemically inducible perIL-15-DmrB was reconstituted in the presence of a small molecule (dimerizer AP20187; +D) but remained inactive in the absence of the dimerizer (-D). (D) The metabolic activity of the NK92 cell line cultured in the presence of perIL-15-DmrB (prepared with or without the dimerizer; +/- D), IL-15, or IL-2. (E, F) To evaluate the effect of perIL-15 on the killing capacity of NK92 cells, the latter were cultured for 48 hours with the perIL-15-DmrB supernatant (+/- dimerizer), IL-15, or IL-2. (G) The NK92 cells cultured in the presence of perIL-15-Dmrb (+dimerizer) successfully infiltrated cancer spheroids and disrupted their architecture. Similar results were achieved using IL-15 or IL-2. In the absence of the dimerizer, IL-15 or IL-2, the NK92 cells infiltrated only a minor portion of the spheroid, mostly remaining attached to the outer layer of the cancer spheroid. BF – bright field. Green – NK92 cells labeled with a Cell Trace Violet stain. Plots show the means ± SEM of at least 9 replicates combined from at least three independent experiments. Conditions were compared using a t-test or one-way ANOVA with a Tukey’s multiple comparisons post-hoc test (\*\*\*\**P* < 0.0001; \*\*\**P*<0.001; \*\**P* < 0.01; \**P*<0.05).

For an external control over the stimulation of immune cells, we developed the chemically inducible perIL-15 by genetically fusing it to the DmrB homodimerization domain. PerIL-15-DmrB stimulated the reporter cell line upon the addition of the small molecule while remaining inactive in the absence of the chemical regulator (Figure 4C). To explore the potential applications of perIL-15-DmrB, we tested the effect of the permuted cytokine on the natural killer NK92 cell line that requires IL-2 or IL-15 supplementation for survival. We show that chemical regulation of the reconstitution of perIL-15-DmrB stimulates the metabolic activity of NK92 cells that reach the same level of metabolic activity as the control cells that were cultured with the standard cytokines IL-15 or IL-2. In contrast, in the absence of the small molecule, the metabolic activity and fitness of NK92 cells were drastically diminished (Figure 4D). To test how the perIL-15 treatment affects the killing capacity, perIL-15-primed (with or without the dimerizer) NK cells were co-cultured with the Raji cancer cells. After 24 hours, the NK92 cells cultured with perIL-15 (+ dimerizer), IL-15, or IL-2 exhibited strongly augmented killing capacity in comparison to the NK92 cells primed with perIL-15 without dimerizer or a cytokine (Figure 4E, F).

To investigate whether the perIL-15 treatment improves the infiltration of NK92 cells into cancer spheroids, we established spheroids of HeLa cells and co-cultured them with NK92 cells that were treated with the perIL-15 supernatant (with or without the dimerizer) or culture medium (with or without the IL-15 or IL-2). After 5 days the NK92 cell treated with the perIL-15 in the presence of the dimerizer disrupted the architecture of the spheroid, resulting in a cell suspension with a few minor cell clumps. Similarly, when the NK92 cells were cultured in the presence of the IL-15 or IL-2, the spheroid architecture was completely obliterated, resulting in a cell suspension. When the NK92 cells were cultured in the presence of perIL-15 without the dimerizer, a minimal infiltration of NK92 cells into the spheroid was detected, and most of the cells were attached to the outer layer of the cancer spheroid. The NK92 cells grown in the absence of the cytokine were not able to infiltrate the cancer spheroid but just attached to the outer layer of the spheroid, while the cancer cells continued to proliferate resulting in an increased size of the spheroid (Figure 4G).

To conclude, our results showcase the potential of using permuted cytokines to improve the anticancer properties of NK cells using a small molecule regulator as a potential safety switch.

## 3 Discussion

Protein-protein interactions govern many cellular processes. In the last two decades, several protein complementation systems have been developed. While the two-component conditional reconstitution systems have been demonstrated also in a therapeutic context (e.g. for anticancer treatment ^32,80–82^ or viral diagnostics^83,84^), their translation is somewhat hindered by the necessity of using two components to reassemble an active protein. Aiming to develop a strategy to enable precise, tunable, and regulated control of cellular processes, we designed the PROPER system, a dimerization-dependent permuted protein platform based on a single polypeptide chain.

A critical step in the design of permuted proteins is the selection of the split site as it determines the scope of their activity. To avoid spontaneous reconstitution, split sites that only enable reconstitution under the induced proximity conditions several computational strategies have been reported (e.g. SPORT^85^, SPELL^9^, Int&in^86^). To prevent spontaneous reconstitution, we not only carefully selected the split site and introduced the permutation but also introduced different types of linkers between the protein domains. The linker could be short and flexible as in the case of perTEVp and nanoluc designs, while an insertion of an extended rigid helix hindered the functionality of a monomeric protein. The strategy of insertion of a rigid linker could be applied for a wide range of proteins, including those with the N- and C-terminus in close proximity, that maintain the activity of a monomer by cyclic permutation. The results on perTEVp designs confirm that the choice of the split site is the most important determinant of the activity of the permuted protein, which could be further fine-tuned by the linkers between the domains of the permuted protein / the permuted protein and a dimerization domain, which primarily need to be sufficiently long to prevent steric hindrance.

Despite conditional reconstitution systems having wide applicability and well-known shortcomings, only a few studies focused on solving the issue of having to deliver two separate constructs to target cells. In 2022, Renna et al. developed a genetically encoded switchable single-chain TEV protease that contained the uniRapR domain, whose conformation is controlled by rapamycin binding^40^. In the absence of a small molecule, the uniRapR domain induces structural disorder resulting in the inactivity of the protein and upon the addition of rapamycin the uniRapR domain and the host protein (TEV protease) become functional. The requirement for the two recombinant proteins by be solved by the delivery of genetically encoded proteins using a 2A system or a polycistronic expression of split proteins separated only by stop codons^35^. Those strategies may however result in an unbalanced production of the two components.

The main advantage of the PROPER system is the reduced complexity, particularly important for therapeutic proteins. The PROPER system requires only a single component that needs to be produced and validated without the need to take care of the precise stoichiometry and possible differences between the two components^87^. Therefore, the development of permuted proteins offers a promising alternative that requires the delivery of a single construct.

We have demonstrated diverse strategies to induce dimerization and reconstruction of the permuted protein functionality, including coiled coil-based dimerization to chemically and intrinsically di/oligomeric biological systems opening various possibilities for novel applications. Furthermore, by demonstrating a permuted protease, a nanoluciferase, and an IL-15 cytokine we demonstrated that this approach can be adopted for proteins of varying folds, sizes, and functions.

To the best of our knowledge, we demonstrate for the first time that the control of cellular processes can be achieved by using a protein complementation system based on a single polypeptide chain. Permuted proteins that can be conditionally reconstituted by di/oligomerization expand the chemical biology toolbox and could be utilized for diverse applications from investigating important biological questions to the development of specific targeted therapies.

## 4 Methods

### 4.1 Design and modeling of permuted proteins

Molecular models of permuted TEV protease, permuted nanoluciferase, LCB3, and permuted IL-15 were generated by Alphafold2^42^ through a locally installed Colabfold^43^ version 1.5.2 using Alphafold multimer weights version 2.3. To create models of permutated TEV and permutated LUC we used Python module Modeller^44,45^ and Chimera^46^. The pdbs of the Alphafold2-modelled components in Chimera were arranged so their relative positions and orientations corresponded to the positions of the predicted fold of the structure. When AF2 was not able to construct a model of permuted dimers, a Modeller was used to connect the segments by specified linkers.

### 4.2 Construct preparation

The perTEV, perNanoLuc, and perIL-15 constructs were purchased as synthetic genes from Twist Bioscience (USA) or IDT (USA) and inserted into the pcDNA3.1 (Invitrogen) vector with Gibson assembly protocol. Plasmids encoding the NS3 protease and CARD8 were gifts from Dustin J Maly (Addgene plasmid number #133616^47^) and Daniel Bachovchin (Addgene plasmid number #169991^48^), respectively. For CARD8 constructs, the amplification of DNA fragments was done using repliQa HiFi ToughMix (Quantabio, USA) and primers purchased from IDT (USA) in PCR reactions performed according to manufacturer instructions. The DNA fragments were inserted into the pcDNA3.1 vector with Gibson assembly protocol, which was carried out using a mixture of the enzymes Taq Ligase (NEB, USA), Phusion Polymerase, and T5 exonuclease (NEB, USA) in reaction buffer (NEB, USA) following the formulation provided by the manufacturer^49^. Plasmid propagation was carried out using the *E. coli* strain DH5-α (NEB, USA). The sequences of all constructs used in this study are provided in Supplemental Table 1.

### 4.3 Mammalian cell culture and transfection

Human kidney epithelial cells (HEK293T), NK-92, and HeLa cells were obtained from the American Type Culture Collection (CRL-11268, CRL-2407, and CCL-2, respectively; ATCC, USA). Raji cells were a kind gift from N. Kopitar-Jerala. The HEK293T and HeLa cells were cultured at 37°C and 5% CO_2_ atmosphere in Dulbecco’s Modified Eagle’s Medium (DMEM) with GlutaMax (Gibco, Life Technologies, USA), supplemented with 10% v/v Fetal Bovine Serum (FBS) (Gibco, Life Technologies, USA). The NK-92 cells were cultured at 37°C and 5% CO_2_ atmosphere in RPMI supplemented with human IL-2 (50 IU/ml) and 20% v/v FBS. The Raji cells were cultured at 37°C and 5% CO_2_ atmosphere in RPMI supplemented with 20% v/v FBS. All cell lines were tested mycoplasma-negative and were authenticated by morphology only.

For luciferase experiments and cell viability testing, 2.0-2.4 × 10^4^ cells per well were seeded in CoStar white or black 96-well plates with an optically clear bottom (Corning, USA). For confocal microscopy experiments, 6.6-7.0 × 10^4^ cells per chamber were seeded in eight-well tissue culture chambers (m-Slide eight-well, Ibidi, Germany). Transfection was performed at 60-70% confluence of HEK293T cells with a mixture of DNA and in-house prepared polyethyleneimine (12 µl per 1000 ng of DNA; stock concentration 0.324 mg/ml, pH 7.5). In the case of chemically-inducible homodimeric (DmrB) or heterodimeric designs (FKBP/FRB), 24 hours after transfection the cells were treated with the AP20187 dimerizer (100 nM for perTEV and 500 nM for peril-15 constructs; MedChemExpress, USA) or rapamycin (100 nM; MedChemExpress, USA), respectively. In the case of HIV inhibitor Ritonavir, the inhibitor was added to cells (100 µM; MedChemExpress, USA) at the time of transfection.

### 4.4 Luciferase assay

To analyze Firefly luciferase activity the HEK293T cells or HeLa cells were lysed 48 hours after transfection using the passive lysis buffer (Biotium, USA). Next, the luciferin substrate (2 mM; Perkin Elmer, USA) supplemented with cofactors (ATP, Sigma, USA; DTT, AppliChem, Germany; coenzyme A, Sigma, USA) was added and the luminescence was measured using a microplate reader (Centro LB 963, Berthold Technologies, Germany).

To analyze the permuted nanoluciferase activity, the HEK293T cells were lysed 48 hours after transfection using the passive lysis buffer (Biotium, USA). To test the activity of permuted nanoluciferase in vitro, the isolated permuted nanoluciferase was added to the HEK293T cells expressing the SARS-CoV-2, SARS-CoV-1, or MERS-CoV spike proteins or mixed directly with the isolated SARS-CoV-2 spike protein. Next, furimazine substrate (final concentration 50 µM; Abmole BioScience, USA) was added and the luminescence was measured using a microplate reader (Centro LB 963, Berthold Technologies, Germany).

### 4.5 Detection of cell membrane permeabilization by propidium iodide uptake

To measure cell membrane permeabilization, propidium iodide was added to the cells at a final concentration of 3.33 µg/ml (Thermo Fisher Scientific, USA), followed by a 30-minute incubation at 37°C and 5% CO_2_ atmosphere. The cells were then centrifuged at 500 × g for 5 minutes and fluorescence was measured using the multiplate reader SynergyMx (BioTek, USA) and Gen 5.1.10 software (BioTek, USA) with program settings bottom reading of fluorescence with an excitation wavelength of 530 nm and emission wavelength of 617 nm. The signal was normalized to untransfected cells as 100% live cells and lysis buffer-treated untransfected cells as 100% dead cells.

### 4.6 Detection of plasma membrane damage by LDH assay

To measure the lactate dehydrogenase release in cell culture supernatants the CyQuant LDH cytotoxicity assay kit (Thermo Fisher Scientific, USA) was used according to the manufacturer’s instructions. Namely, 50 µl of cell culture supernatants were transferred to a new 96-well plate and mixed with 50 µl of LDH assay buffer. After 15-20 minutes of incubation at room temperature, the absorbance was measured at 492 nm and 680 nm using a multiplate reader SinergyMx (BioTek, USA) and Gen 5.1.10 software (BioTek, USA). The signal was normalized to the supernatant of untransfected cells as a negative control and the supernatant of lysis buffer-treated untransfected cells as a positive control.

### 4.7 Confocal microscopy

48 hours after transfection the cells were stained with Hoechst 33342 (1 μg/ml; Thermo Fisher Scientific, USA) and 7-AAD viability dye (1.25 μg/ml; eBioscience, Invitrogen, Thermo Fisher Scientific, USA) or propidium iodide (3.33 µg/ml; Thermo Fisher Scientific, USA) for 10-15 minutes at 37°C. The samples were imaged using a Leica TCS SP5 inverted laser-scanning microscope on a Leica DMI 6000 CS module equipped with an HCX PL Fluotar L ×20, numerical aperture 0.4 (Leica Microsystems, Germany). Image processing was performed with Leica LAS AF Lite software (Leica Microsystems, Germany).

### 4.8 Protein expression of permuted nanoluciferase protein

The expression of permuted nanoluciferase fused to LCB3 was performed in bacteria using *E. coli* strain NiCO21(DE3) (NEB, MA USA) transformed with the expression plasmid. We prepared the preculture by inoculating 100 ml LB media supplemented with Kanamycin (50 μg/ml) and incubated it at 37 °C, 160 RPM overnight. The next day the culture was grown at 37 °C until reaching OD 0.6 - 0.9. At that point, we induced the protein expression by the addition of 1 mM IPTG (Goldbio, USA) and grew the culture overnight in agitation (180 RPM) at 20 °C. After centrifugation, the bacterial pellets were resuspended in 40 ml of lysis buffer 50 mM Tris–HCl at pH 8.0, 150 mM NaCl, 10 mM imidazole, 18 U/ml Benzonase (Merck, Germany), 1 mM MgCl_2_, 2 μl/ml CPI (Protease Inhibitor Cocktails) (Millex Sigma-Aldrich, USA) per liter of culture. Lysis was done by ultrasonication with a Vibra-cell VCX (Sonics, USA) on wet ice for a maximum of three cycles of 2 min of total pulse time, at intervals of 1 s pulse and 3 s pause (60% amplitude). The cellular lysates were centrifuged at 16000×g (4 °C) for 20 min and afterward, the soluble fraction was filtered through 0.45 μm filter units (Sartorius Stedim, Germany) and stored for further purification.

### 4.9 Production of recombinant viral proteins

Recombinant viral proteins were produced as previously described^50^. Briefly, the prefusion ectodomain of the SARS-CoV-2 spike protein was first transiently transfected into suspension-adapted Expi293 cells (Thermo Fisher Scientific, USA) using the PEI MAX transfection reagent (Polysciences, USA) and a ProCHO5 culture medium (Lonza, Switzerland). One hour following transfection the dimethyl sulfoxide (DMSO; AppliChem, Germany) was supplemented to a final concentration 2% (v/v,) and the cells were incubated on a shaker for five days at 31 °C and 4.5% CO2. Next, the Strep-Tactin column (IBA Lifesciences, Germany) was used to purify the clarified supernatant, followed by dialysis into a PBS buffer.

### 4.10 Protein chromatography

Protein isolation of permuted nanoluciferase fused to LCB3 consisted of two chromatography steps: affinity (Ni-NTA) and size exclusion chromatography (SEC). The filtered bacterial lysates were incubated with 5 ml of Ni-NTA resin (Goldbio, USA) equilibrated with buffer A (50 mM Tris–HCl pH 8.0, 150 mM NaCl, 10 mM imidazole) for 5 minutes. The resin was washed with 200 ml of buffer A and 200 ml of buffer B (50 mM Tris–HCl pH 8.0, 150 mM NaCl, 20 mM imidazole). Afterward, we performed elution of the bound protein with buffer C (50 mM Tris–HCl pH 8.0, 150 mM NaCl, 300 mM imidazole). The eluted protein was further purified with SEC using 320 ml of HiLoad Superdex™ 75 resin (GE Healthcare, USA) packed in a 26/600 XK column (GE Healthcare, IL USA) equilibrated with filtered and degassed SEC buffer (20 mM Tris–HCl pH 7.5, 150 mM NaCl, 10% v/v glycerol). 12 ml of NiNTA elution was filtered through 0.22 μm syringe filters (Millex Sigma-Aldrich, USA) before being injected into the column. The chromatography was run using an AKTA™ pure FPLC system (GE Healthcare, IL USA) in SEC buffer at a linear flow rate of 2.6 ml/min and the eluted protein fractions of 4 ml were collected separately.

### 4.11 Size exclusion chromatography coupled with multi-angle light scattering

We performed size exclusion chromatography multi-angle light scattering (SEC-MALS) measurements using an HPLC system (Waters, USA) coupled with a UV detector, a Dawn8+ multiple-angle light scattering detector (Wyatt, USA) and a refractive index detector (RI500, Shodex, Japan). We filtered the protein sample through 0.1 μm Durapore centrifuge filters (Merck Millipore, USA) before being injected onto a Superdex 200 increase 10/300 column (GE Healthcare, USA) equilibrated with SEC buffer B (20 mM Tris–HCl pH 7.5, 150 mM NaCl). The data analysis was performed using Astra 7.0 software (Wyatt, USA), and the peak of interest was analyzed.

### 4.12 perIL-15 binding and activity measurements

To test the activity of wild-type interleukin (IL) 15 and perIL-15 variants, the HEK-Blue CD122/CD132 reporter cell line that expresses an inducible secreted embryonic alkaline phosphatase (SEAP) after human IL-15 receptor activation was used (Invivogen, USA). Reporter cells were maintained in DMEM (Gibco, Life Technologies, USA) supplemented with 4.5 g/L glucose, GlutaMax, and 10% v/v heat-inactivated FBS (Gibco, Life Technologies, USA). To test the activity of perIL-15 constructs, the HEK293T cells were first transfected and the supernatant was collected 48 hours after transfection. Next, 5 × 10^4^ reporter cells per well were seeded in transparent 96-well tissue culture plates (TPP, Switzerland) together with 20 µl of cell culture supernatants containing perIL-15 variants or IL-15. After 24 hours, 20 µl of supernatant was transferred to a new plate and the content of alkaline phosphatase was analyzed based on the change in QUANTI-Blue (Invivogen, USA) absorbance at 630 nm.

### 4.13 Analysis of the effect of permuted IL-15 on immune cells

First, we prepared the supernatant containing perIL-15 by transiently transfecting the HEK293T cells cultured in RPMI medium (supplemented with 10% FBS) with the perIL15-DmrB construct using PEI. After 24 hours the cells were treated with 500 nM AP20187 and 48 hours later the supernatant was collected and stored at -20°C. To assess the effect of perIL-15 on the metabolism of NK-92 cells, 1×10^5^ cells per well were seeded on a black 96-well plate (Costar) in a culture medium containing perIL-15 or culture medium alone (both supplemented with 20% FBS). After 72 hours the resazurin was added (50 µM) and after a 4-hour incubation at 37°C and 5% CO_2_, the fluorescence was measured using the multiplate reader SynergyMx (BioTek, USA) and Gen 5.1.10 software (BioTek, USA) with program settings bottom reading of fluorescence with an excitation wavelength of 560 nm and emission wavelength of 590 nm.

To evaluate the effect of perIL-15 on the killing capacity of NK-92 cells we first cultured the NK-92 cells (3×10^5^ cells per well on a 48-well plate, TPP, Switzerland) in the culture medium containing perIL-15 for 48 hours and then added 6×10^4^ Raji cells labeled with CellTrace Far Red (Invitrogen, Thermo Fisher Scientific, USA) per well. After 24 hours the samples were imaged using the CellInsight CX7 LZR Pro HCS platform and cancer cells were quantified based on the intensity of the fluorescence signal that was analyzed using the ImageJ software.

To evaluate the killing capacity of perIL-15-primed NK-92 cells in a 3D setting, we first plated 5×10^3^ of HeLa cells per well on an ultra-low attachment 96-well plate (Corning) in DMEM medium (supplemented with 10% FBS) and NK92 cells in a 6 well plate in RPMI or perIL-15 supernatant (with or without the dimerizer) supplemented with 20% FBS. After 48h in culture, we exchanged the culture medium of spheroids for culture medium containing perIL-15 or culture medium alone and seeded 5×10^4^ of NK92 cells labeled with CellTrace Violet (Invitrogen, Thermo Fisher Scientific, USA) per well. After 24/ 48h and 5 days, the samples were imaged using the CellInsight CX7 LZR Pro HCS platform. Before the last imaging, the culture medium was pipetted up and down three times without disrupting the spheroid.

### 4.14 Statistical analysis

Data are presented as means ± SEM of pooled data from at least three independent experiments. Statistical analyses were performed in the GraphPad Prism 8 software (GraphPad Software, version 8.4.3; USA). The one-way ANOVA with Tukey’s multiple comparisons test was used to analyze data with multiple groups and independent variables.

## ACKNOWLEDGMENTS

This work was funded by Slovenian Research and Innovation Agency project grants Z3-4501 (T.Z.R.), N3-0358 and J3-60056 (I.H.-B), J7-4640 (R.J.), programme grant P4-0176 (R.J.), European Research Council (ERC) Advanced Grant project MaCChines grant agreement ID: 899259 (R.J.) and European Union’s Horizon 2020 Research and Innovation Programme within the FET Open Project Virofight (Grant Agreement No 899619), and EU funding (no. 101059842, CTGCT) (R.J.). The authors thank Petra Dekleva for cloning the permuted nanoluciferase plasmids and Dr. Hana Esih for providing the SARS-CoV-2 spike protein. The authors are grateful to Dr. Tina Fink for advice regarding the stimulation of NK cells.

## Supplemental Figures

**Supplemental Figure 1.**
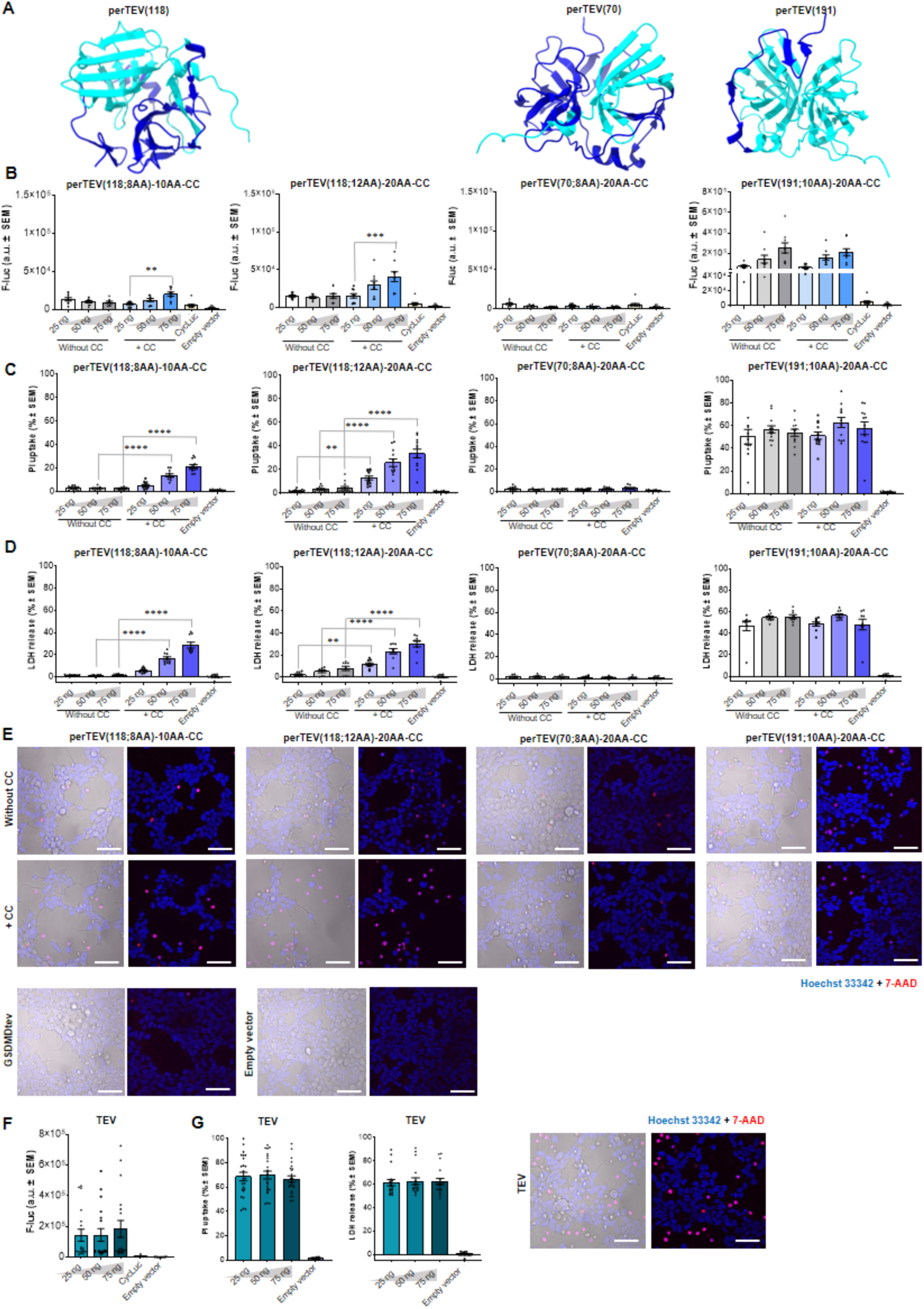
Additional perTEV protease designs. (A) Protein models of perTEV protease with a split at site 118 AA / 70 AA / 191 AA. (B) Using a CycLuc reporter, the activity of additional perTEVp design was tested. The perTEVp designs with a split at site 118AA induced a dose-dependent response when fused to a dimerization domain. The design with a split at site 70AA was inactive and the design with a split at site 191AA was auto-active. (C-E) The perTEV protease designs were used to induce cell death in HEK293T cells co-transfected with GSDMDtev. (F) Using the CycLuc reporter the activity of wild-type TEV protease was assessed. (G) The wild-type TEV protease successfully induced cell death in HEK293T cells co-transfected with the GSDMDtev as demonstrated by PI uptake, LDH release, and confocal microscopy. Scale bars 50 μm. Plots show the means ± SEM of at least 9 replicates combined from at least three independent experiments. Conditions were compared using a one-way ANOVA with a Tukey’s multiple comparisons post-hoc test (\*\*\*\**P* < 0.0001; \*\*\**P*<0.001; \*\**P* < 0.01; \**P*<0.05).

**Supplemental Figure 2.**
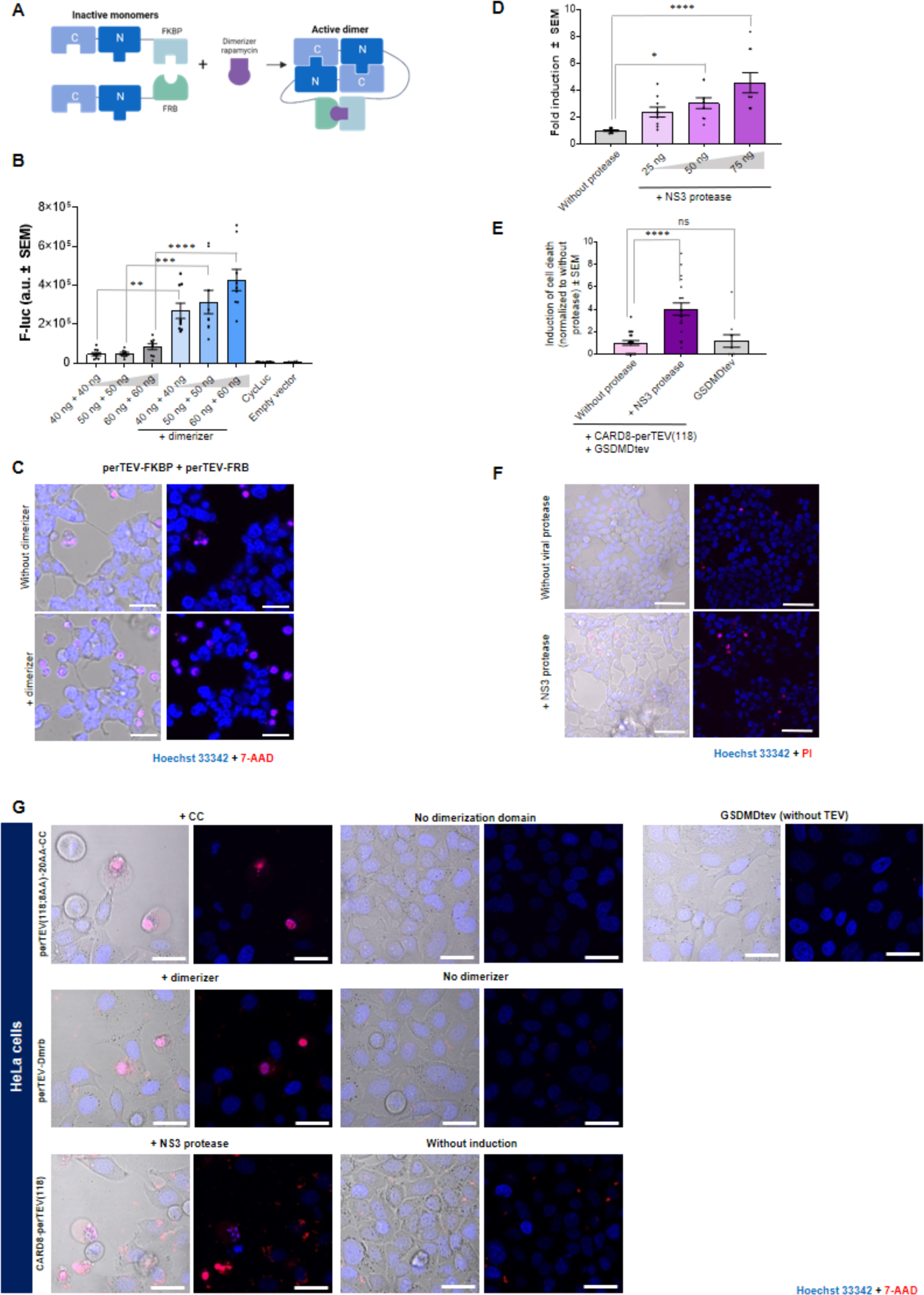
Chemical and biological signals induce reconstitution of perTEV protease that leads to cell death in HEK293T and HeLa cells. (A, B) To demonstrate that chemically inducible dimerization domains can be applied, we prepared perTEV proteases fused to FKBP or FRB domains. Following the addition of a small molecule (rapamycin), the perTEVp reconstituted and successfully cleaved the CycLuc reporter. (C) The perTEVp fused to the FKBP/FRB heterodimerization system can induce cell death in HEK293T cells when co-transfected with gasdermin D with a TEVp cleavage site. (D-F) To develop a biologically inducible system the perTEVp(118; 8AA) was fused to the CARD8. As the CARD8 was previously genetically modified it carried the NS3 protease cleavage site. Following viral cleavage, the CARD8-based effector oligomerized which resulted in the reconstruction of the perTEVp which then cleaved the CycLuc reporter in a dose-dependent manner. (G) To demonstrate that perTEVp designs can induce cell death in cancer cells, the HeLa cells were transiently transfected with GSDMDtev and perTEVp(118;8AA)-20AA-CC/ perTEVp-DmrB /CARD8-perTEVp. All three designs successfully induced cell death in cancer cells following the induction of dimerization. Confocal microscopy images show pyroptotic cells exhibiting typical pyroptotic morphology. Dead cells were stained with PI (red) and nuclei were labeled with Hoechst 33342 (blue). Scale bars (C, G) 20 μm, (F) 50 μm. Plots show the means ± SEM of at least 9 replicates combined from at least three independent experiments. Conditions were compared using a one-way ANOVA with a Tukey’s multiple comparisons post-hoc test (\*\*\*\**P* < 0.0001; \*\*\**P*<0.001; \*\**P* < 0.01; \**P*<0.05).

**Supplemental Figure 3.**
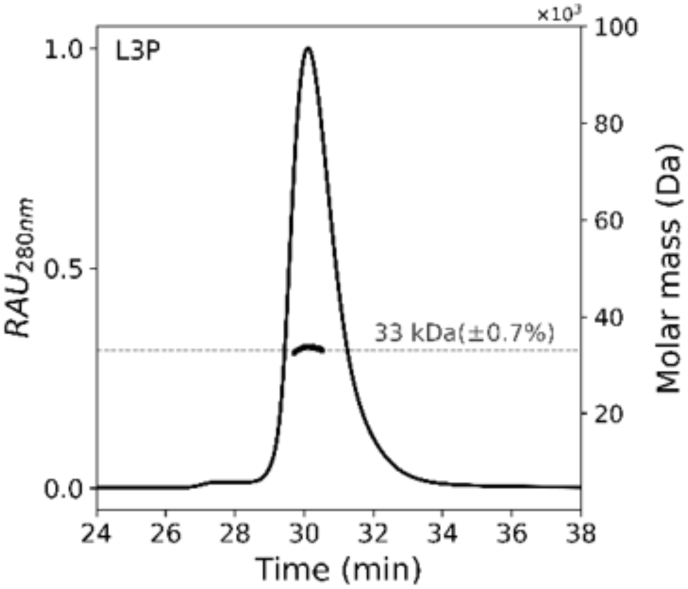
SEC-MALS chromatogram of the purified protein perNanoLuc-L3P. The molecular weight of the peak was calculated from the light scattering and it corresponds to the theoretical mass calculated from the amino acid sequence (theoretical Mw of L3P = 28.6 kDa).

